## Supplementary figures and images for "Hepatic MIR20B promotes nonalcoholic fatty liver disease by suppressing *PPARA*"

### Figure 1-figure supplement 1

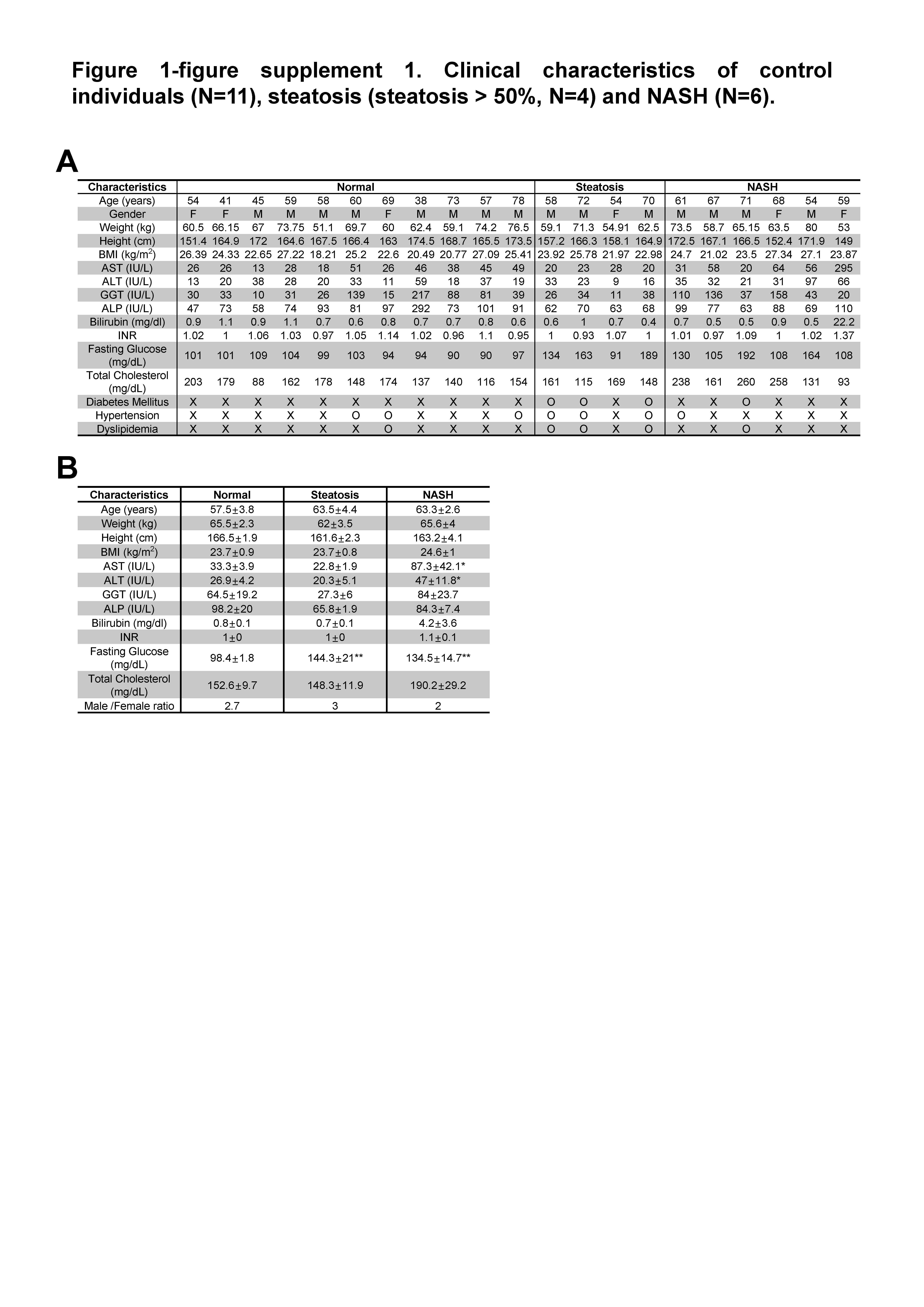

### Figure 1-figure supplement 2

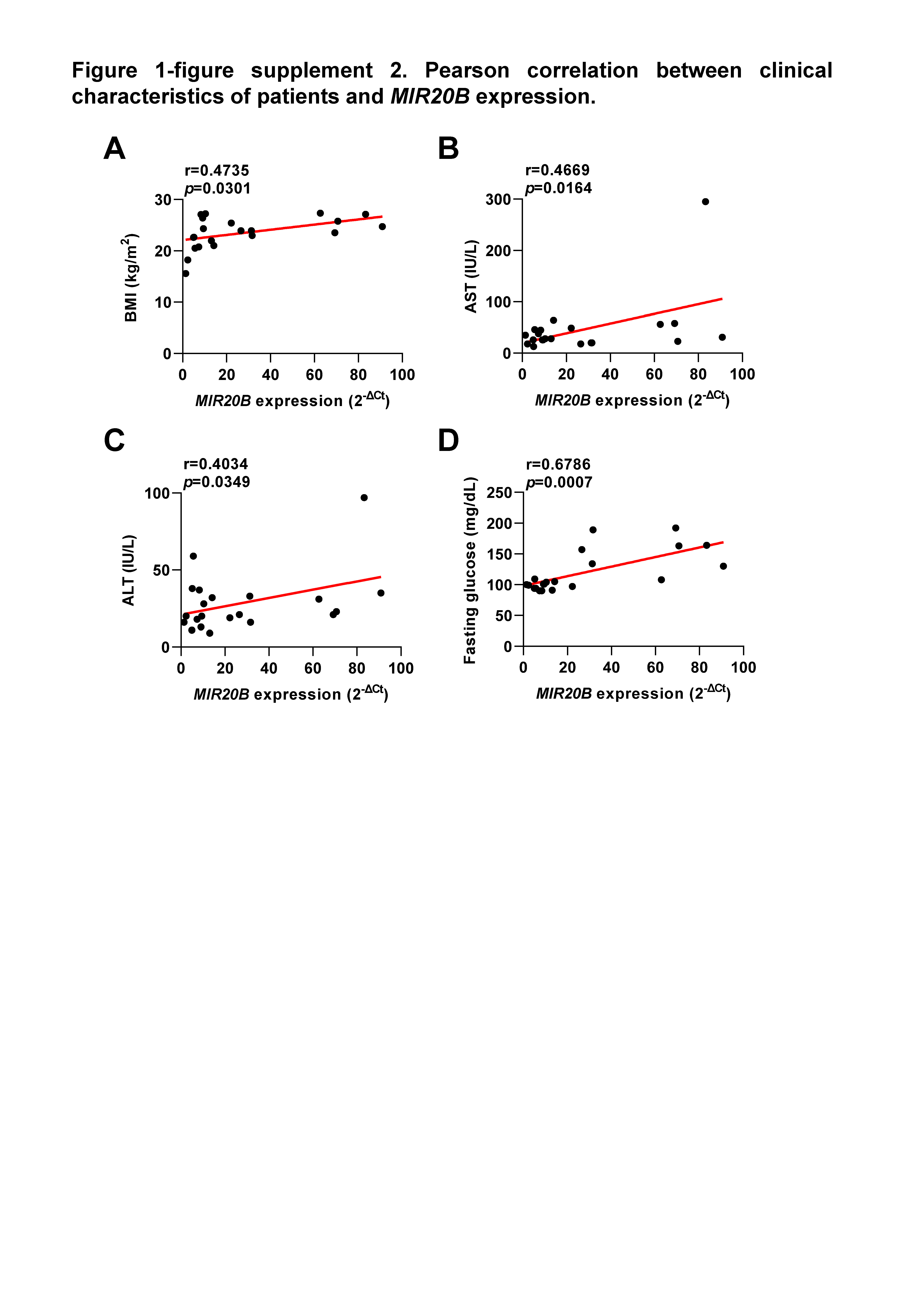

### Figure 1-figure supplement 3

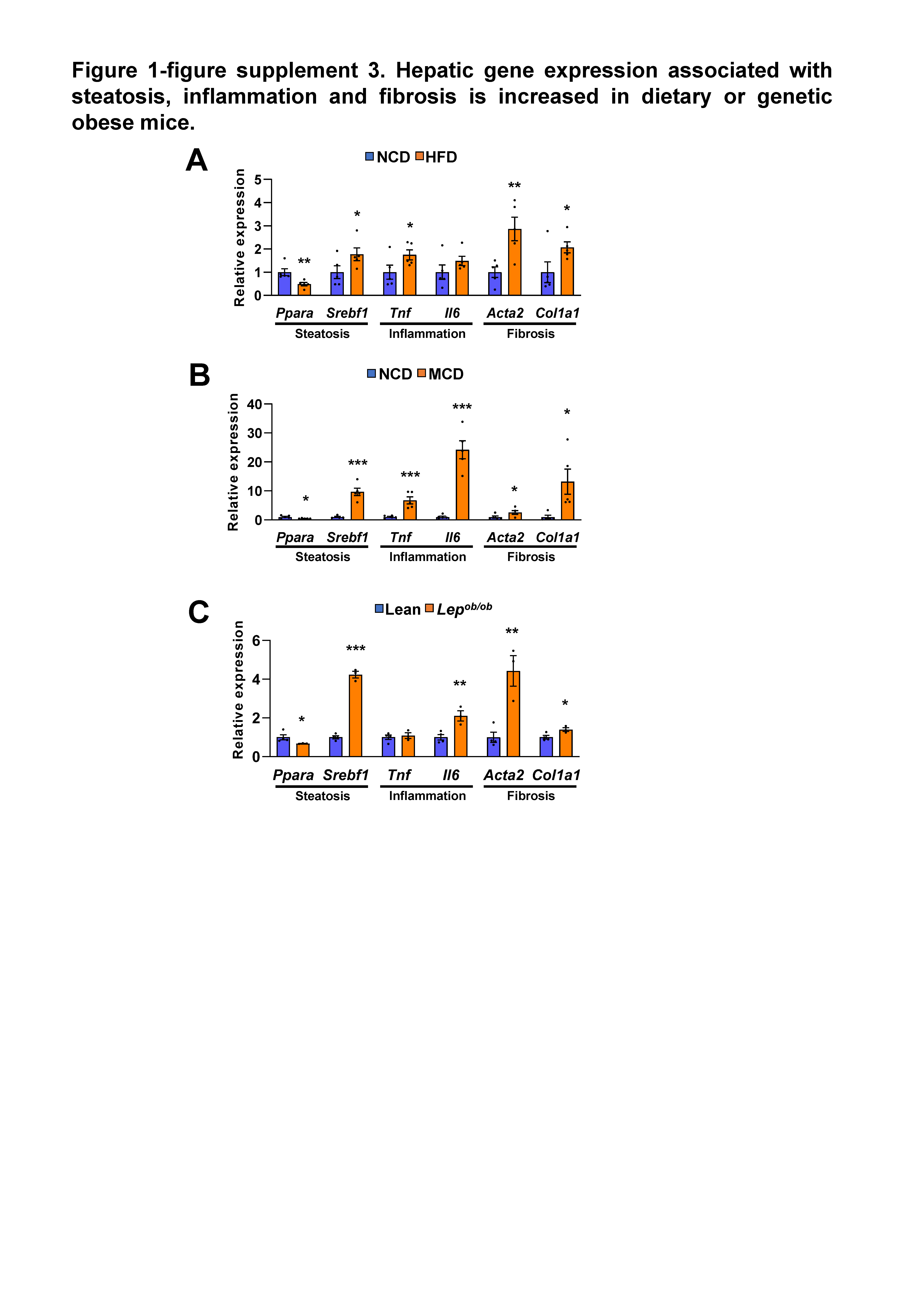

### Figure 1-figure supplement 4

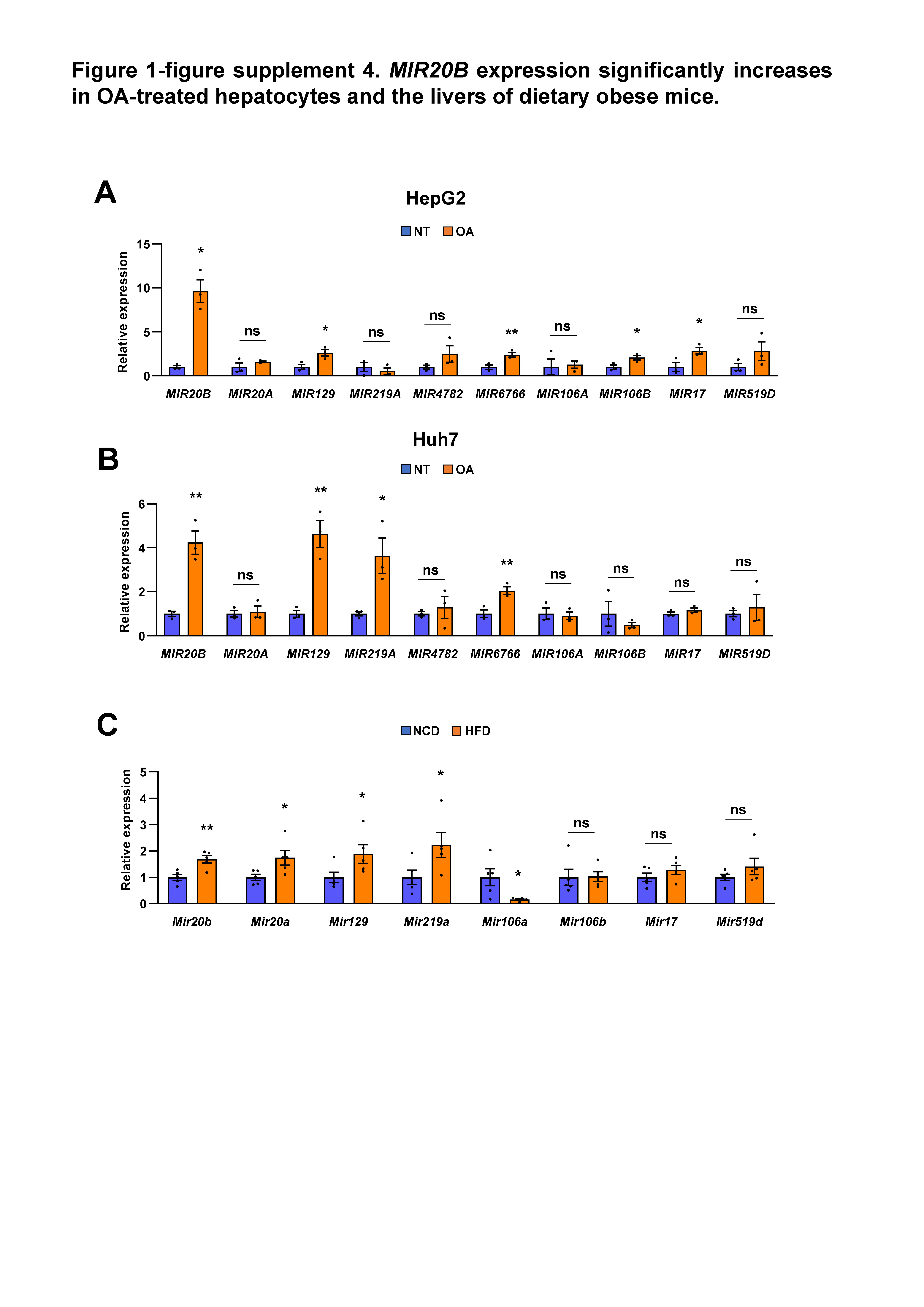

### Figure 2-figure supplement 1

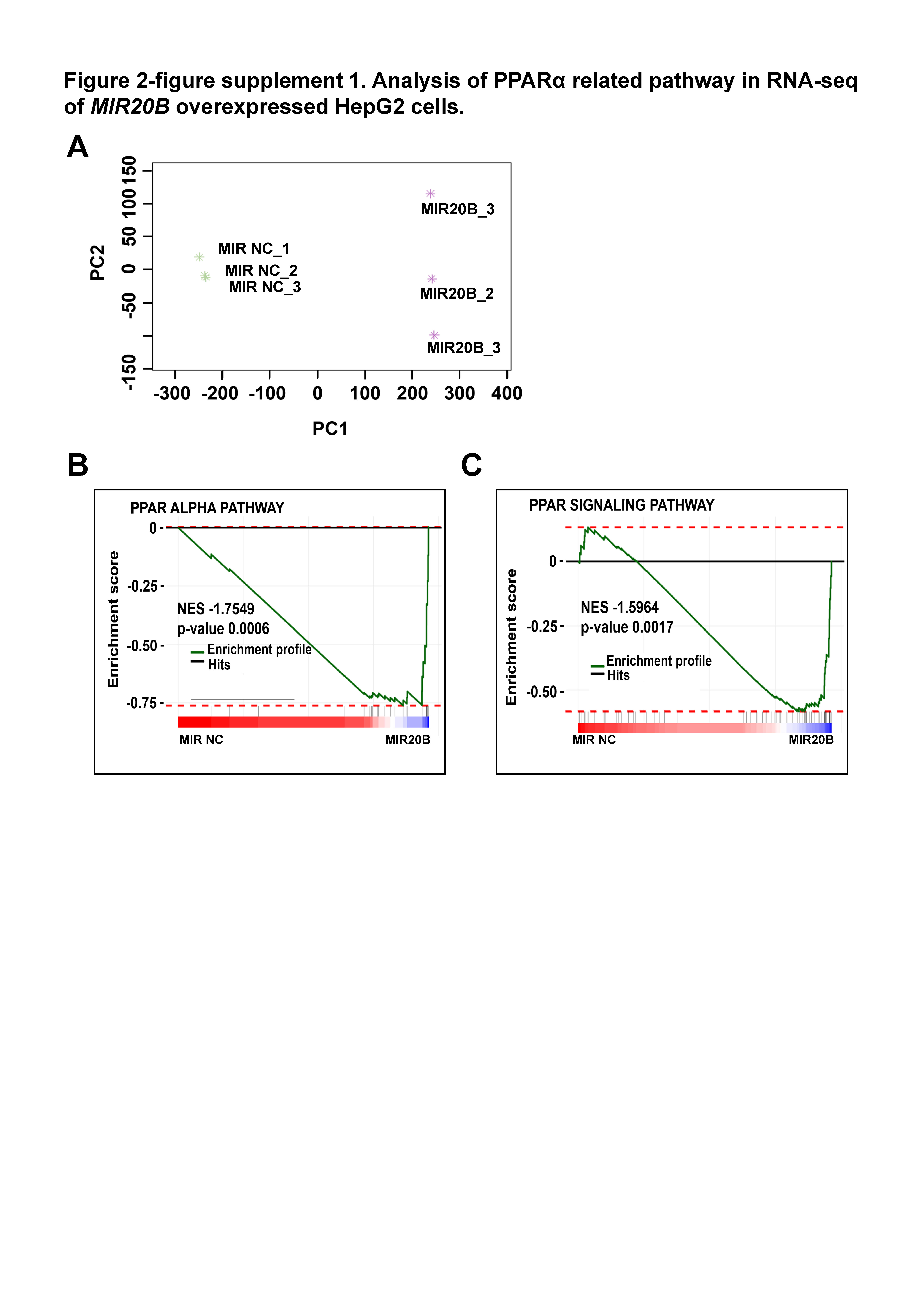

### Figure 2-figure supplement 2

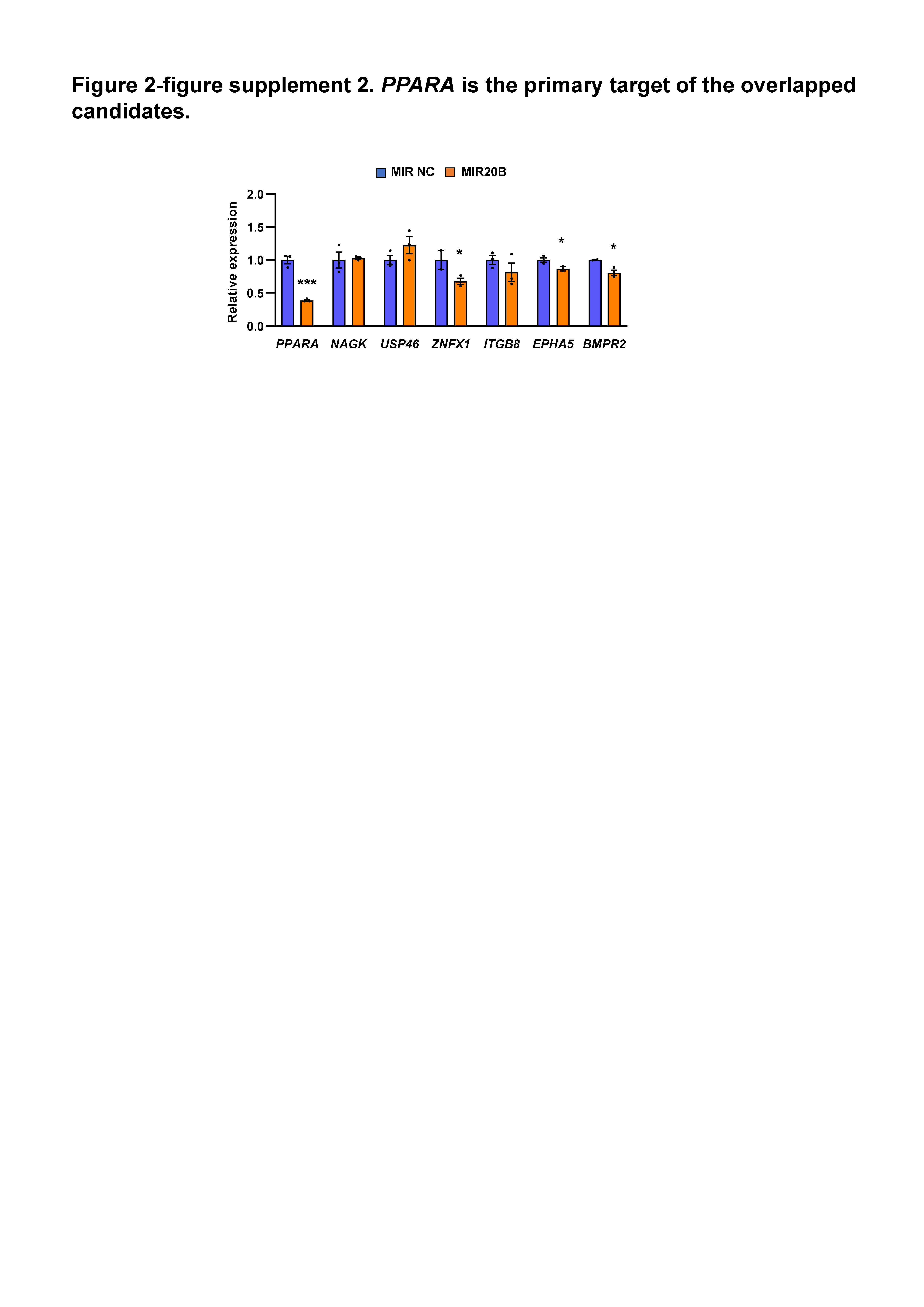

### Figure 2-figure supplement 3

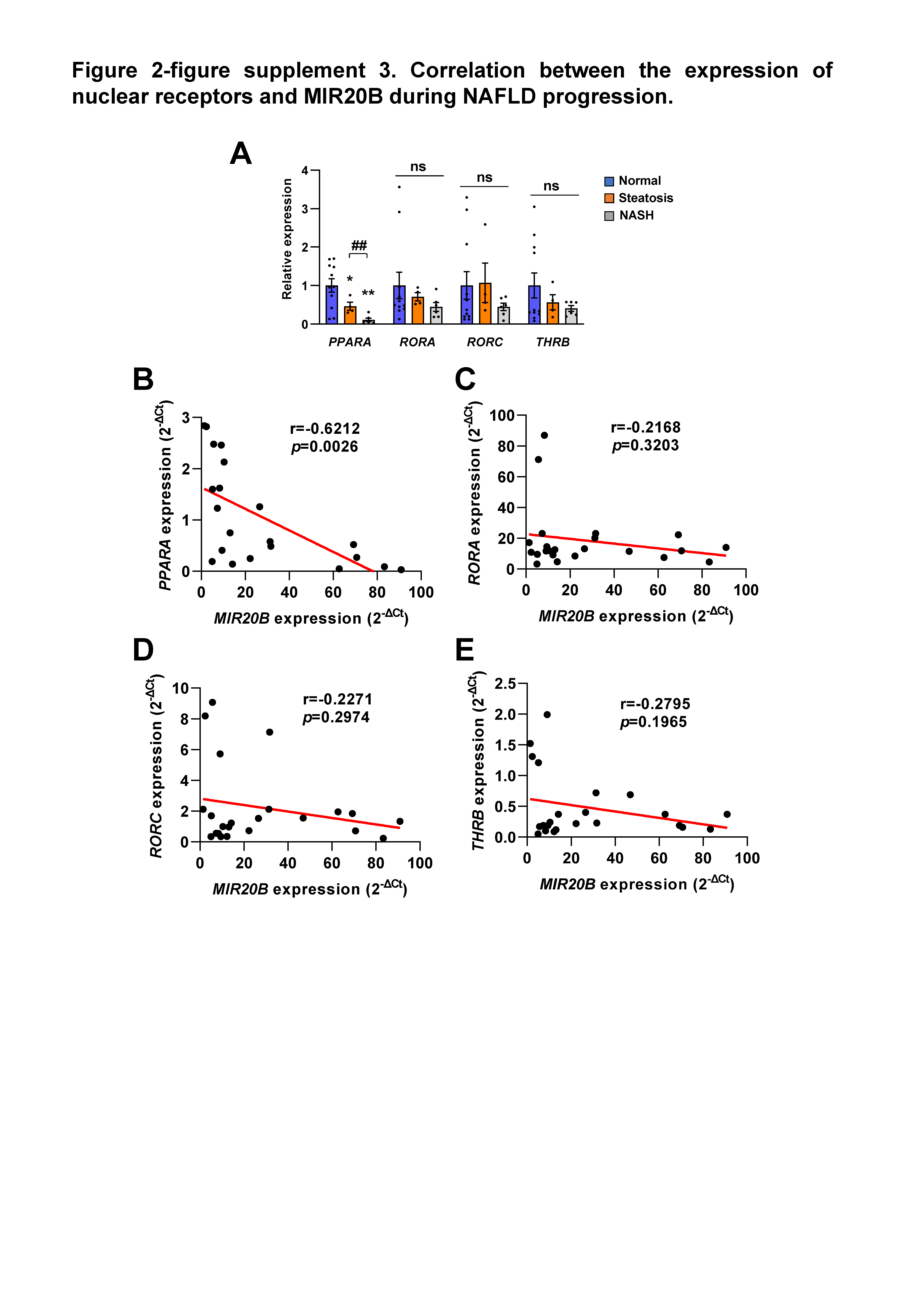

### Figure 2-figure supplement 4

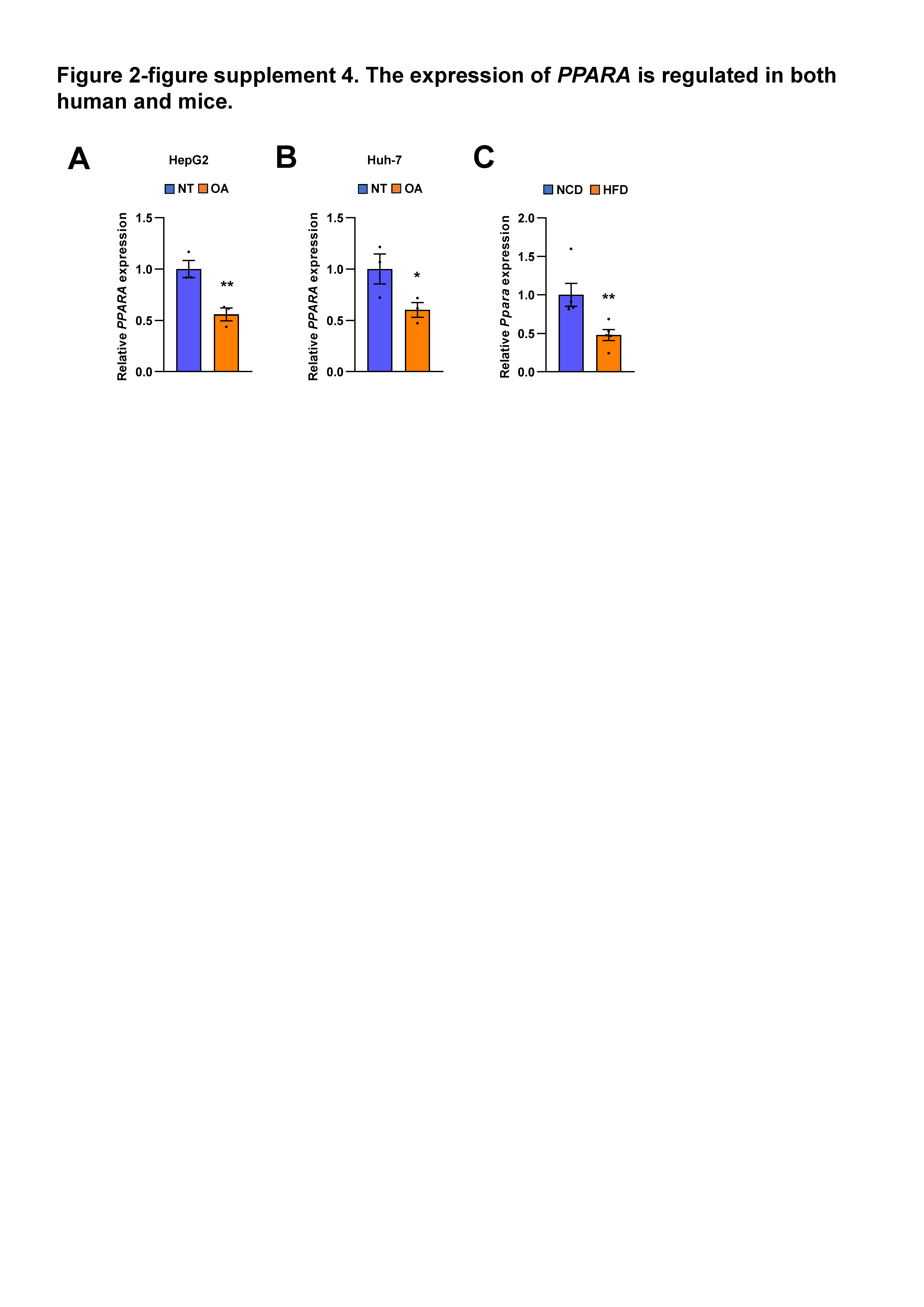

### Figure 2-figure supplement 5

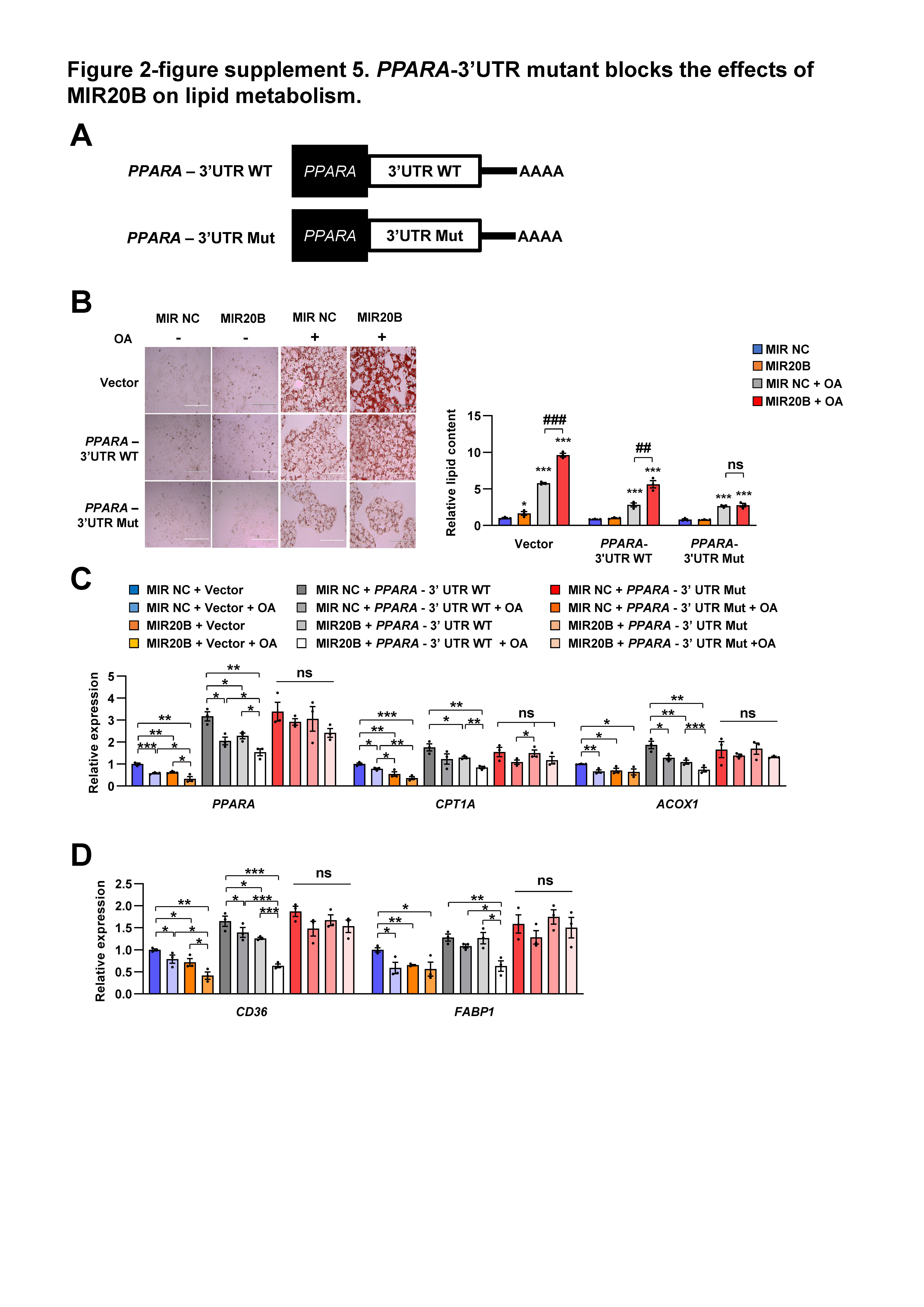

### Figure 4-figure supplement 1

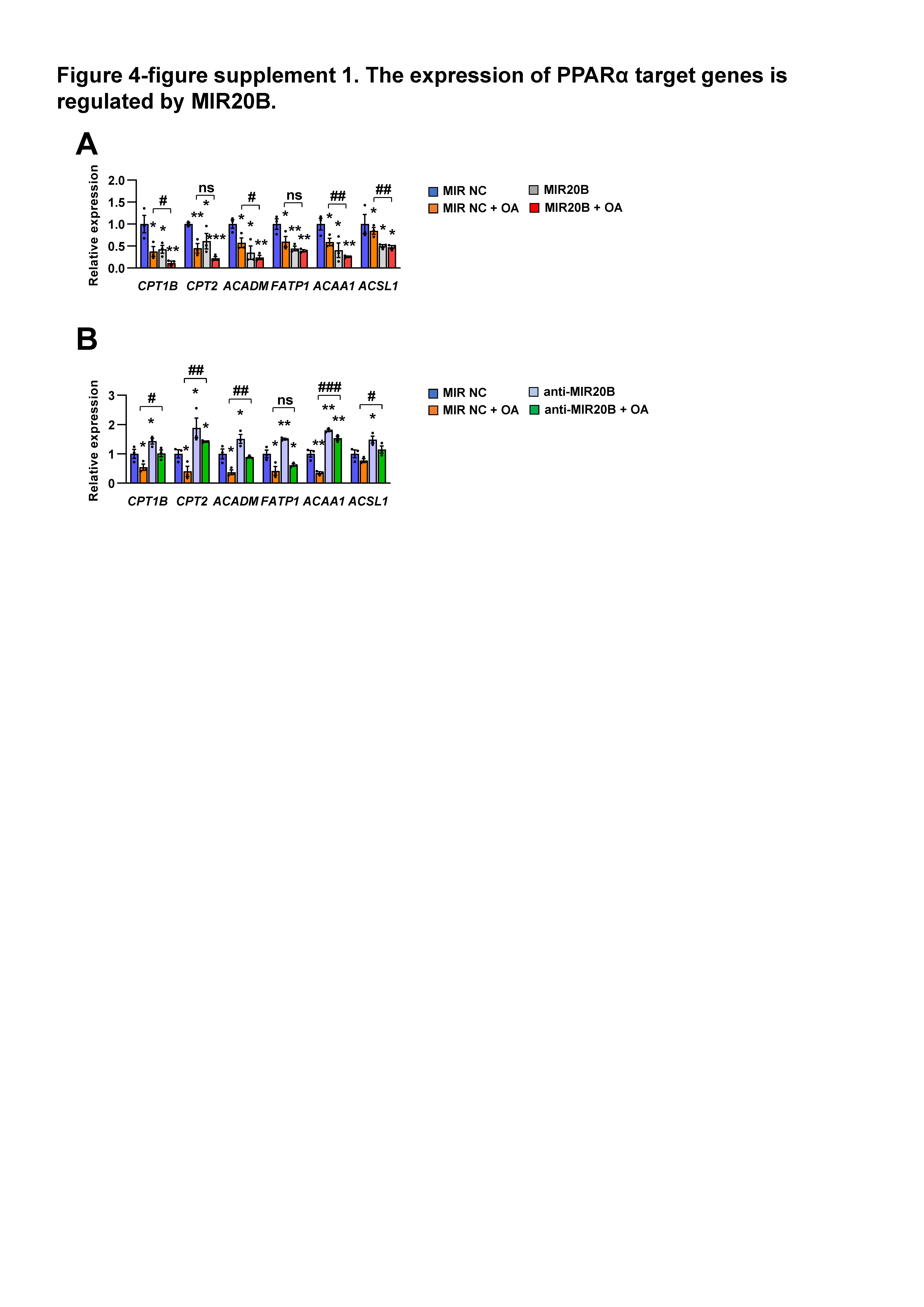

### Figure 4-figure supplement 2

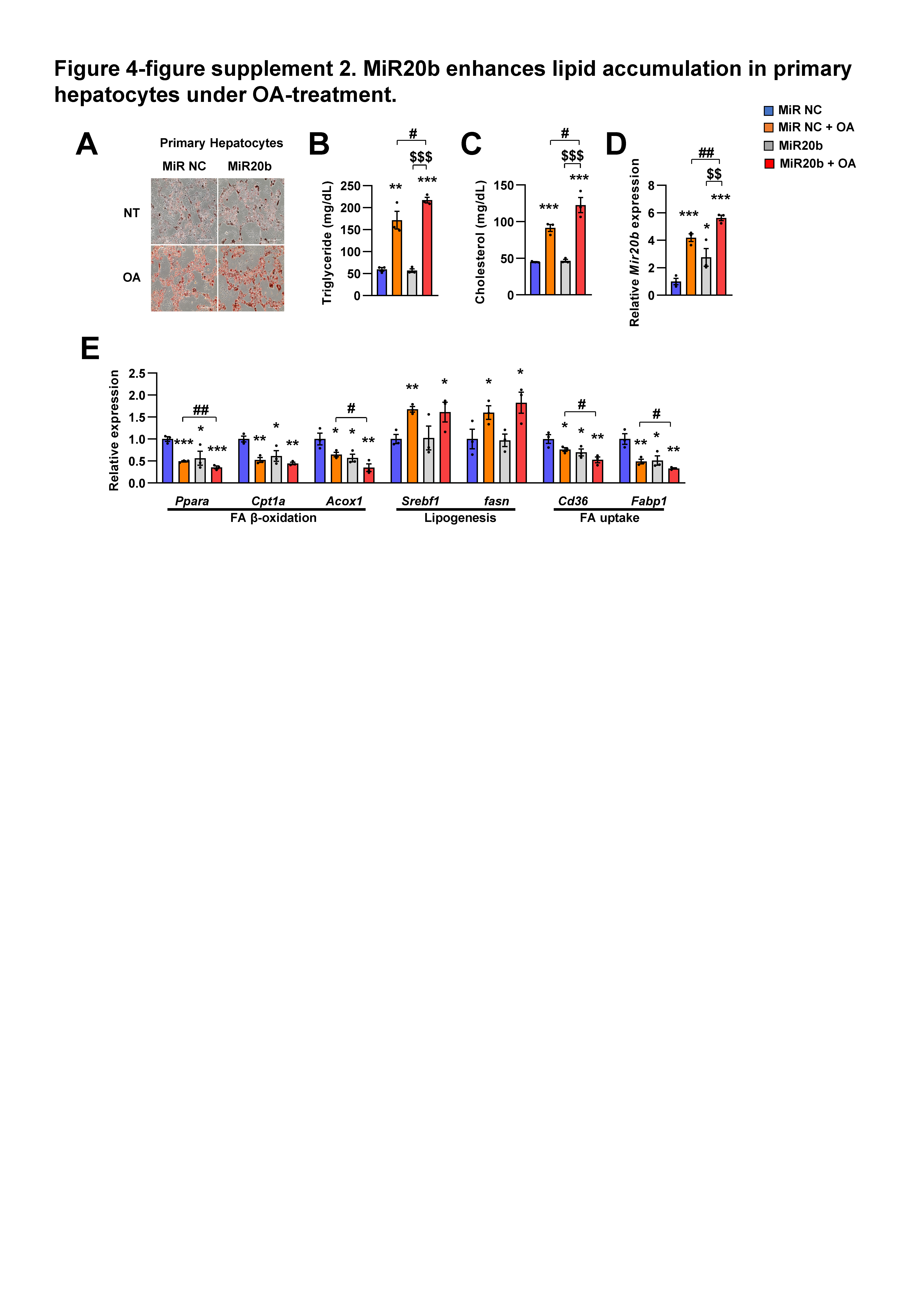

### Figure 4-figure supplement 3

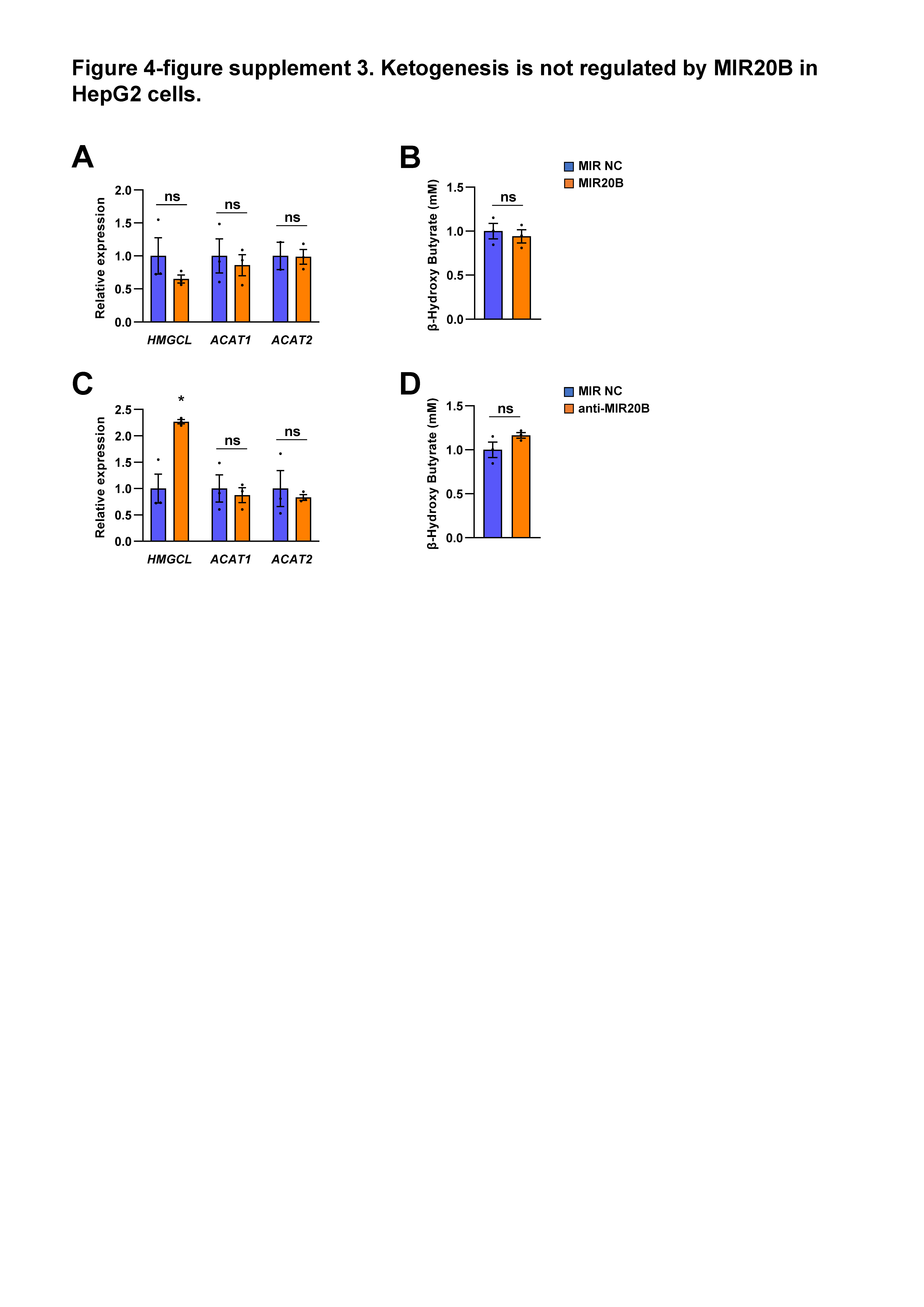

### Figure 4-figure supplement 4

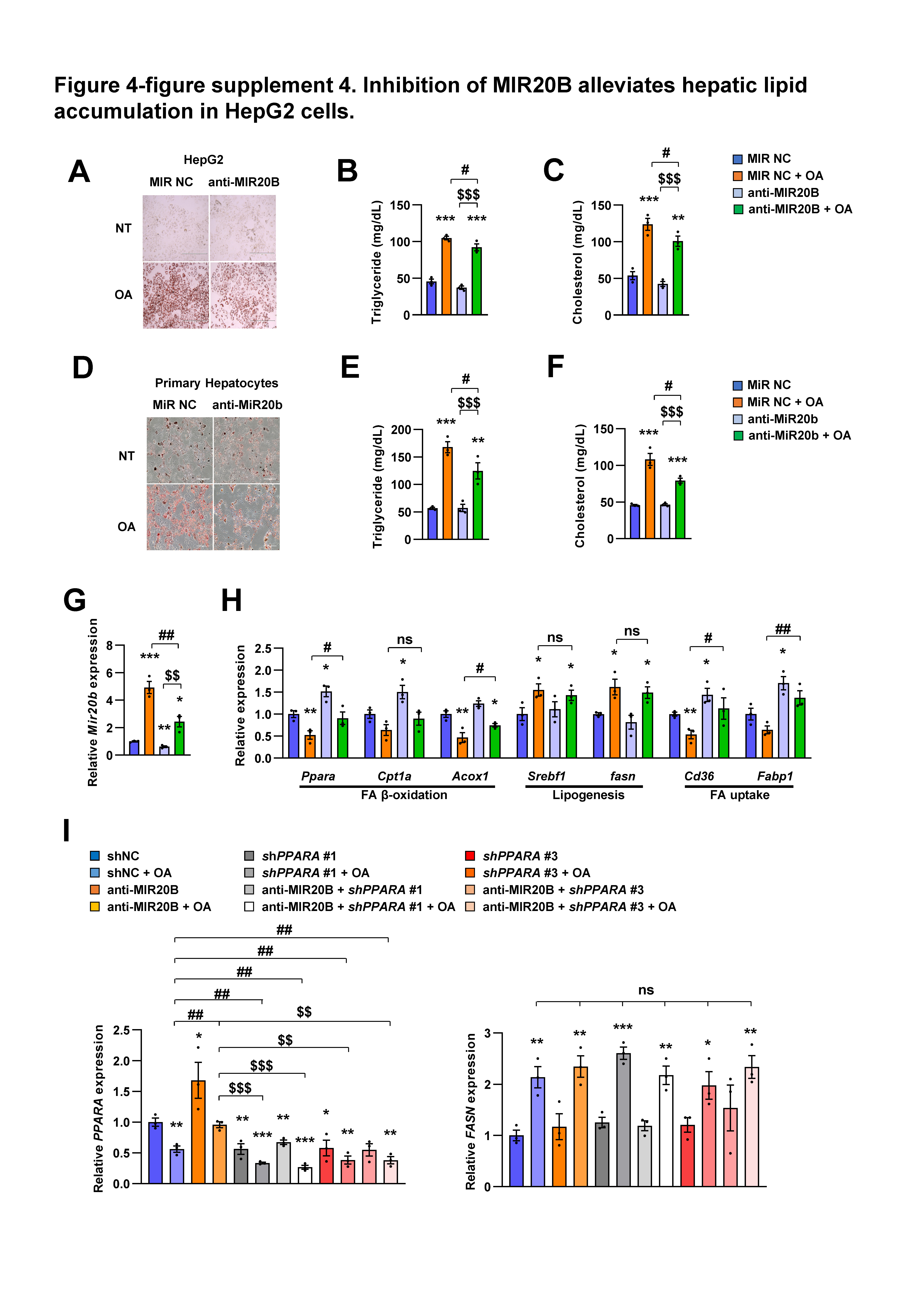

### Figure 4-figure supplement 5

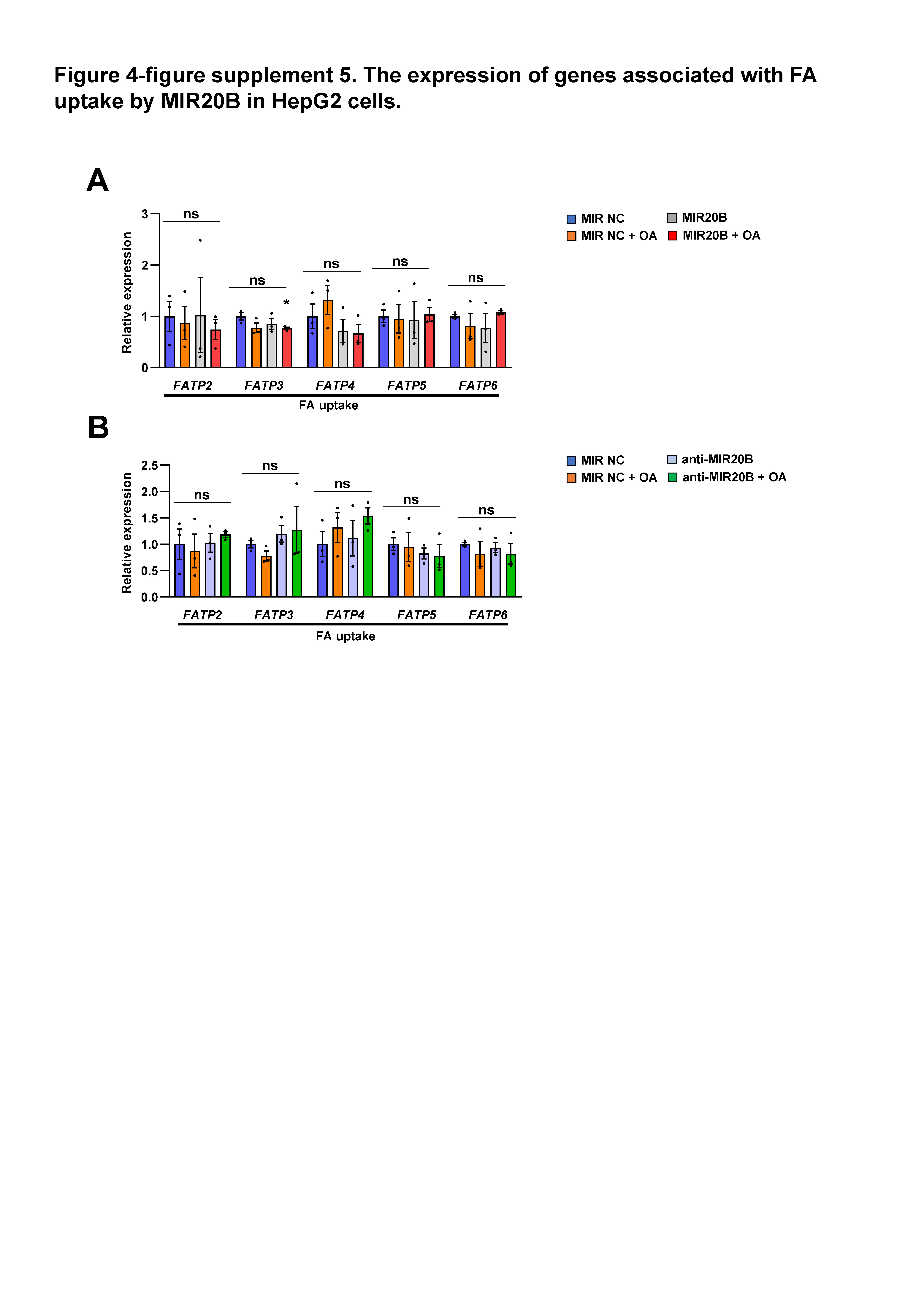

### Figure 4-figure supplement 6

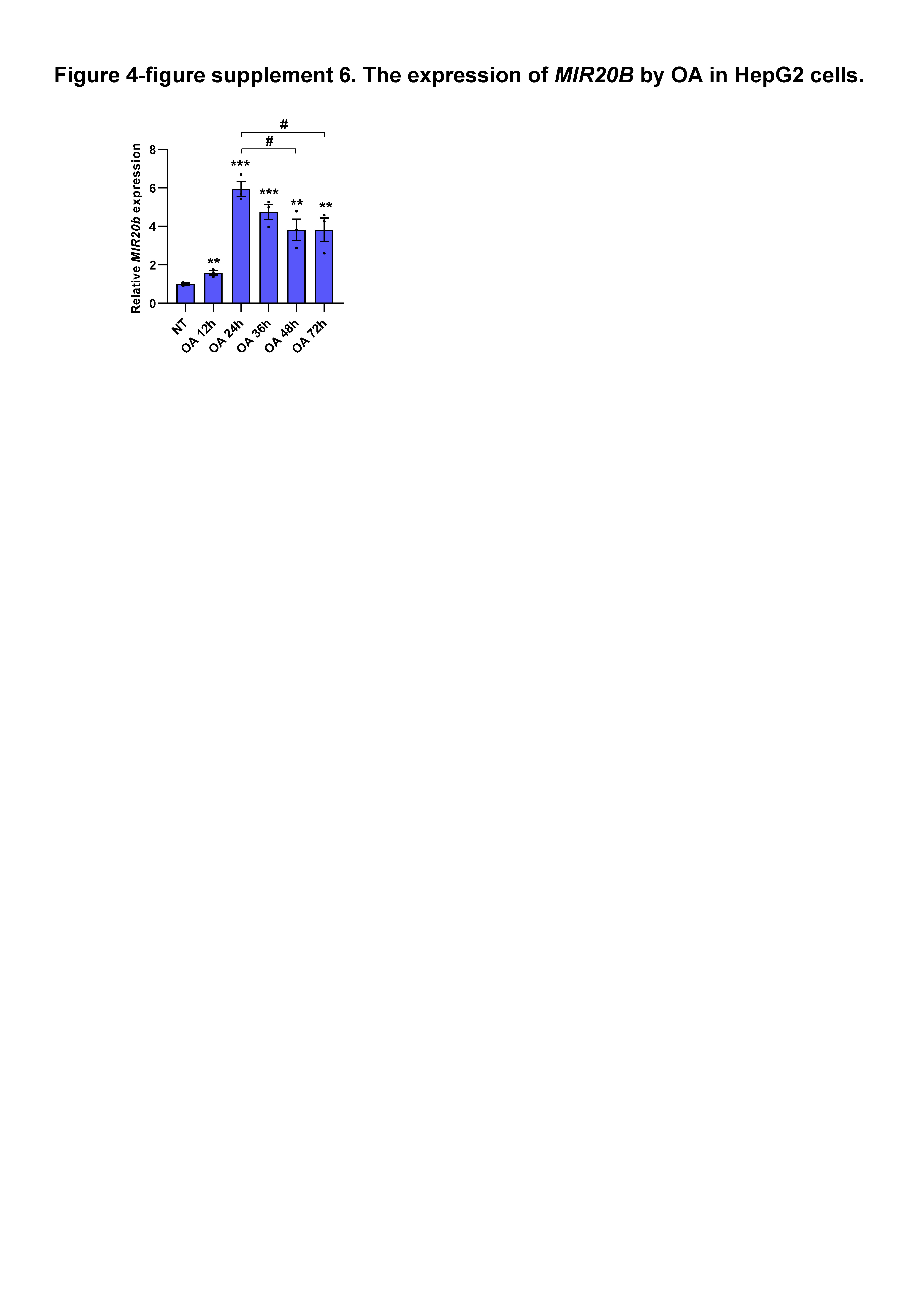

### Figure 5-figure supplement 1

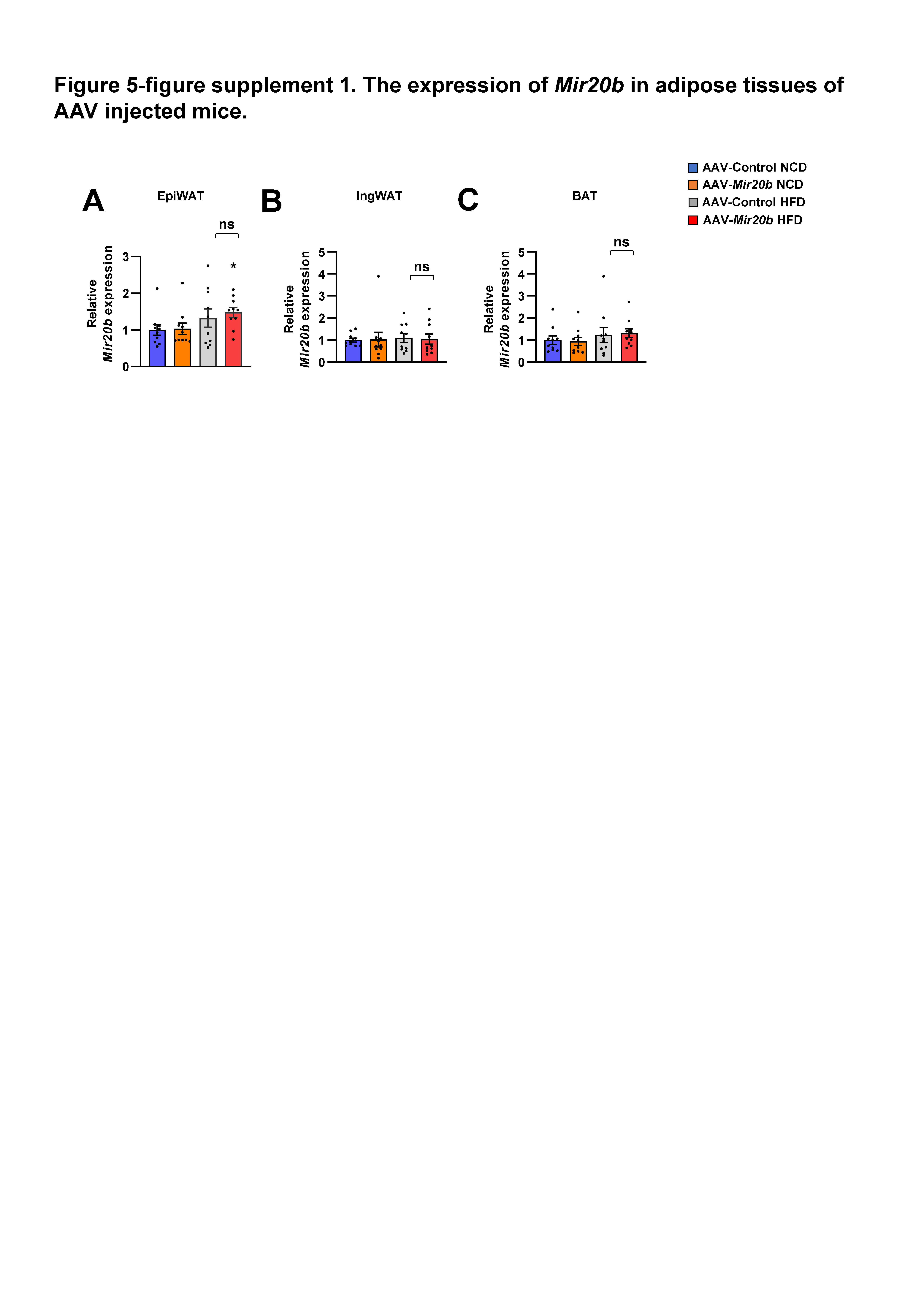

### Figure 5-figure supplement 2

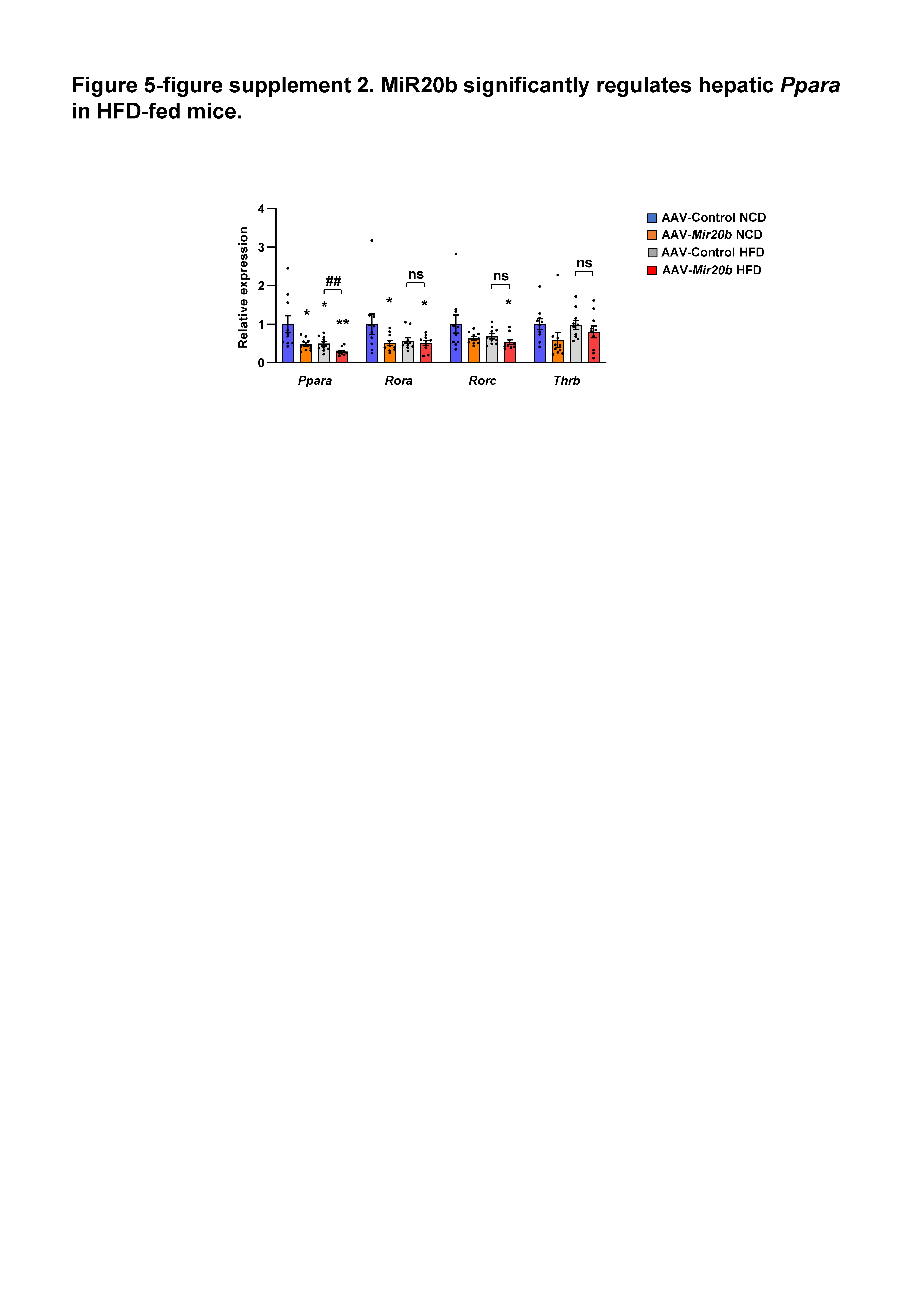

### Figure 5-figure supplement 3

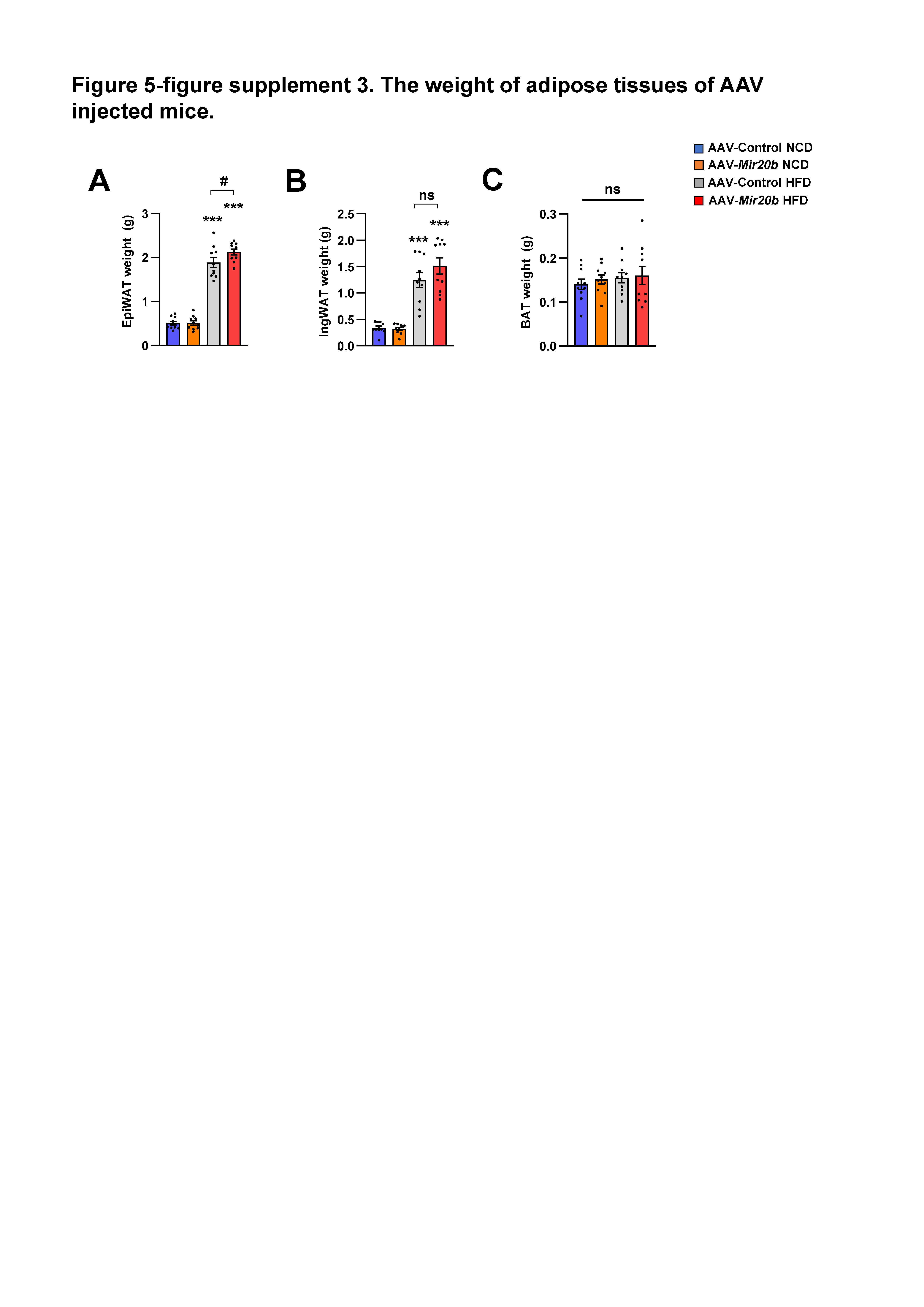

### Figure 5-figure supplement 4

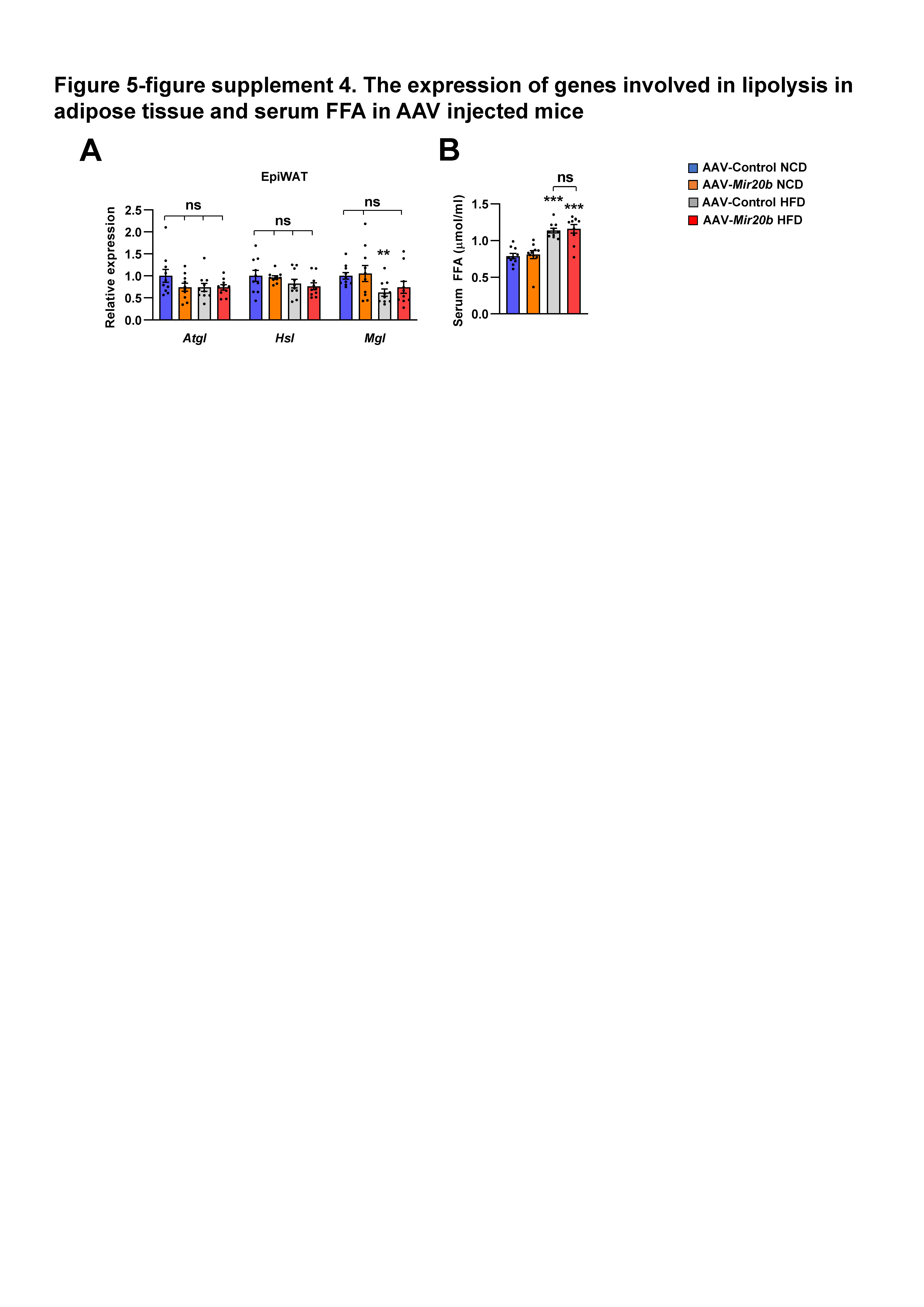

### Figure 6-figure supplement 1

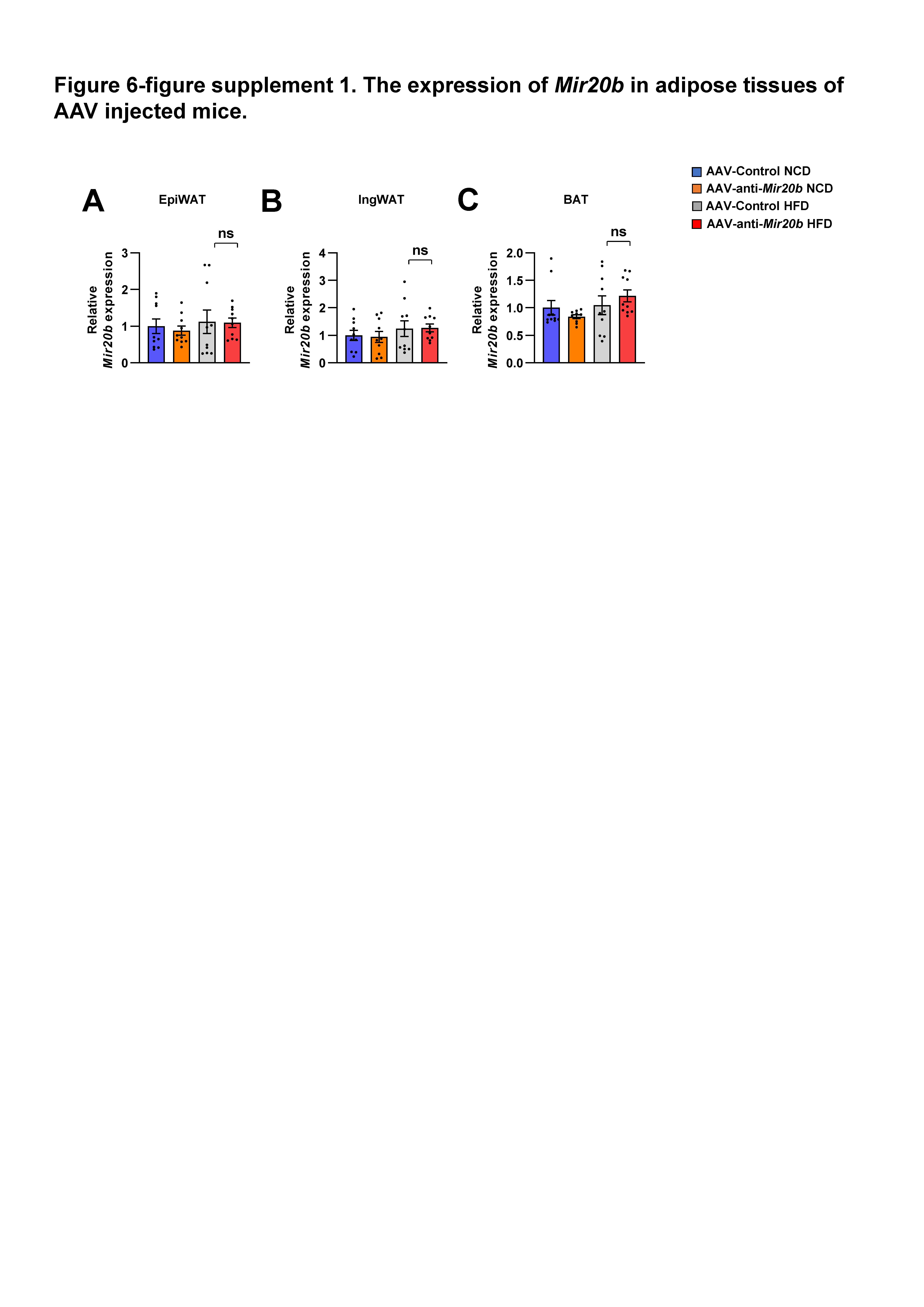

### Figure 6-figure supplement 2

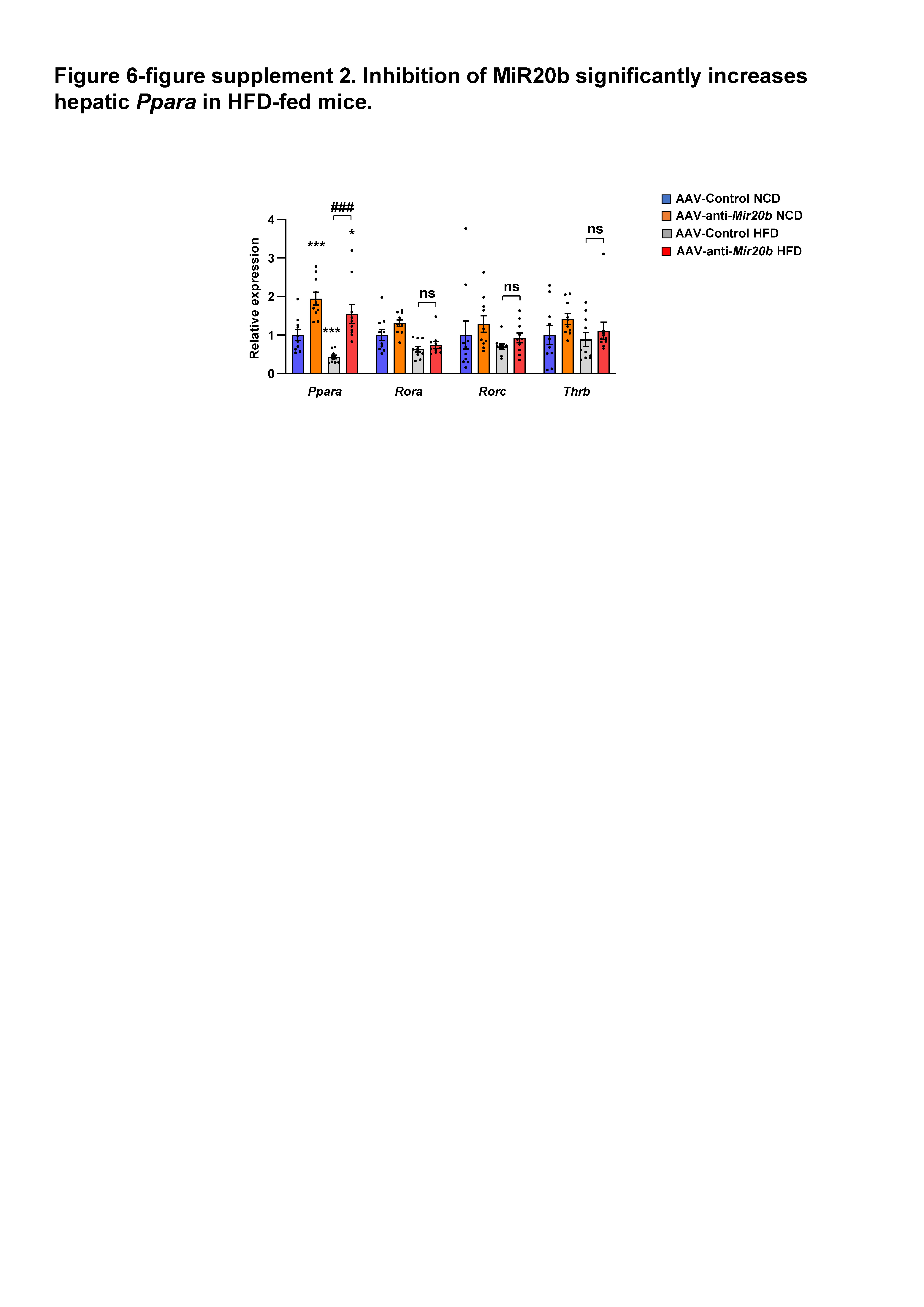

### Figure 6-figure supplement 3

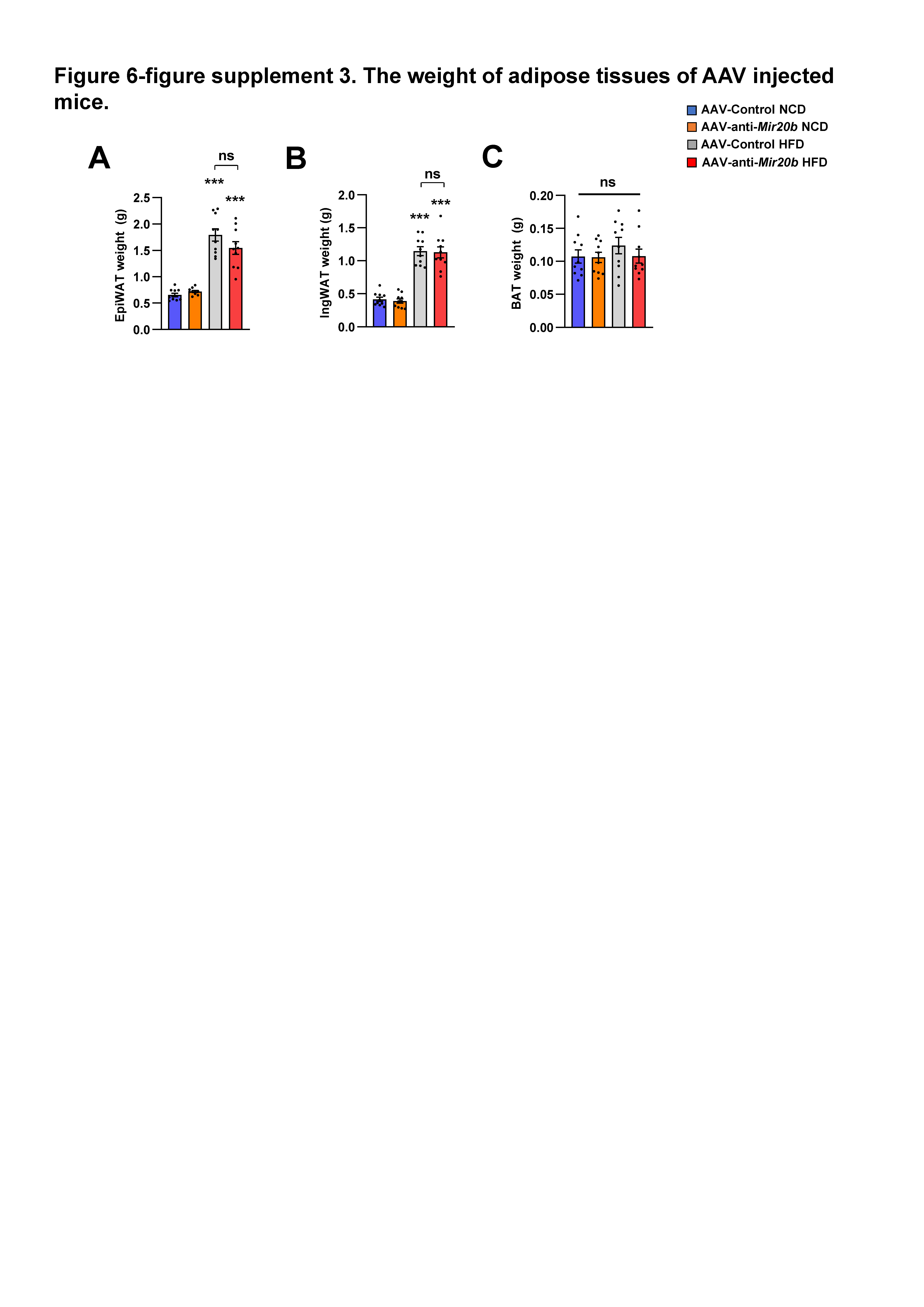

### Figure 6-figure supplement 4

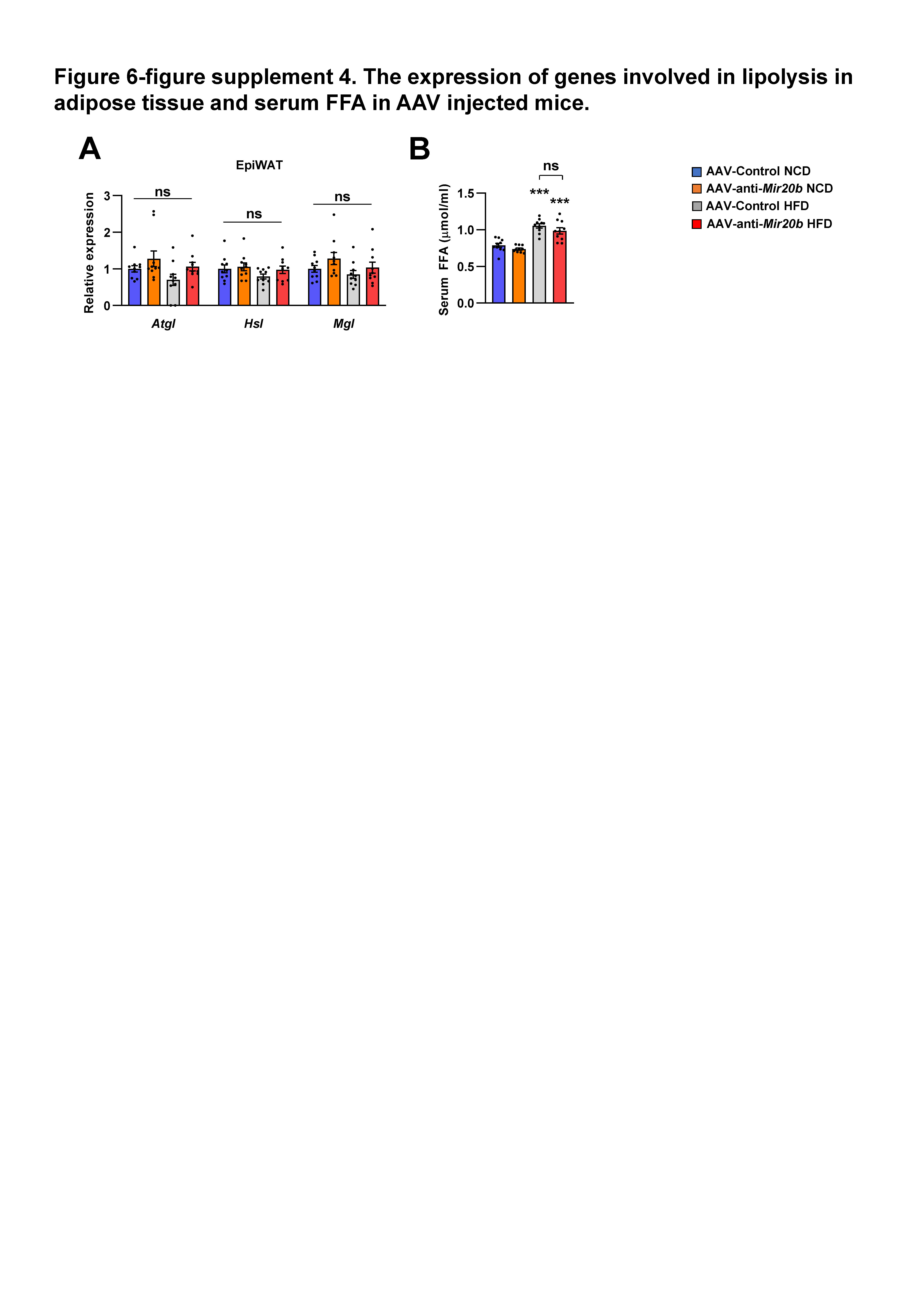
